# Identification of DAXX As A Restriction Factor Of SARS-CoV-2 Through A CRISPR/Cas9 Screen

**DOI:** 10.1101/2021.05.06.442916

**Authors:** Alice Mac Kain, Ghizlane Maarifi, Sophie-Marie Aicher, Nathalie Arhel, Artem Baidaliuk, Sandie Munier, Flora Donati, Thomas Vallet, Quang Dinh Tran, Alexandra Hardy, Maxime Chazal, Françoise Porrot, Molly OhAinle, Jared Carlson-Stevermer, Jennifer Oki, Kevin Holden, Etienne Simon-Lorière, Timothée Bruel, Olivier Schwartz, Sylvie van der Werf, Nolwenn Jouvenet, Sébastien Nisole, Marco Vignuzzi, Ferdinand Roesch

## Abstract

Interferon restricts SARS-CoV-2 replication in cell culture, but only a handful of Interferon Stimulated Genes with antiviral activity against SARS-CoV-2 have been identified. Here, we describe a functional CRISPR/Cas9 screen aiming at identifying SARS-CoV-2 restriction factors. We identified DAXX, a scaffold protein residing in PML nuclear bodies known to limit the replication of DNA viruses and retroviruses, as a potent inhibitor of SARS-CoV-2 and SARS-CoV replication in human cells. Basal expression of DAXX was sufficient to limit the replication of SARS-CoV-2, and DAXX over-expression further restricted infection. In contrast with most of its previously described antiviral activities, DAXX-mediated restriction of SARS-CoV-2 was independent of the SUMOylation pathway. SARS-CoV-2 infection triggered the re-localization of DAXX to cytoplasmic sites and promoted its degradation. Mechanistically, this process was mediated by the viral papain-like protease (PLpro) and the proteasome. Together, these results demonstrate that DAXX restricts SARS-CoV-2, which in turn has evolved a mechanism to counteract its action.

## Introduction

Severe Acute Respiratory Syndrome Coronavirus 2 (SARS-CoV-2) is the causative agent of COVID-19 and the third coronavirus to cause severe disease in humans after the emergence of SARS-CoV in 2002 and Middle East Respiratory Syndrome-related Coronavirus (MERS-CoV) in 2012. Since the beginning of the pandemic, SARS-CoV-2 has infected more than 200 million people and claimed more than 4 million lives. While the majority of infected individuals experience mild (or no) symptoms, severe forms of COVID-19 are associated with respiratory failure, shock and pneumonia. Innate immune responses play a key role in COVID-19 pathogenesis: immune exhaustion (1) and reduced levels of type-I and type-III interferons (IFN) have been observed in the plasma of severe COVID-19 patients (2,3). Imbalanced immune responses to SARS-CoV-2, with a low and delayed IFN response coupled to early and elevated levels of inflammation, have been proposed to be a major driver of COVID-19 (4,5). Neutralizing auto-antibodies against type-I IFN (6) and genetic alterations in several IFN pathway genes (7) have also been detected in critically ill COVID-19 patients. These studies highlight the crucial need to characterize the molecular mechanisms by which IFN effectors may succeed, or fail, to control SARS-CoV-2 infection.

Although SARS-CoV-2 has been described to antagonize the IFN pathway by different mechanisms involving the viral proteins ORF3b, ORF9b ORF6, and Nsp15 (8), detection of SARS-CoV-2 by the innate immune sensor MDA5 (9,10) leads to the synthesis of IFN and expression of IFN Stimulated Genes (ISGs) in human airway epithelial cells (4). IFN strongly inhibits SARS-CoV-2 replication when added in cell culture prior to infection (11,12) or when administered intranasally in hamsters (13), suggesting that some ISGs might have antiviral activity (14). Relatively few ISGs with antiviral activity against SARS-CoV-2, however, have been identified so far. For instance, spike-mediated viral entry and fusion is restricted by LY6E (15,16) and IFITMs (17,18). Mucins have also been suggested to restrict viral entry (19). ZAP, which targets CpG dinucleotides in RNA viruses, also restricts SARS-CoV-2, albeit moderately (20). OAS1 has been recently identified in an ISG overexpression screen to restrict SARS-CoV-2 replication, through the action of RNAseL, both in cell lines and in patients (21). Another overexpression screen identified 65 ISGs as potential inhibitors of SARS-CoV-2 (22), and found that BST-2/Tetherin is able to restrict viral budding, although this activity is counteracted by the viral protein ORF7a. We hypothesize that additional ISGs with antiviral activity against SARS-CoV-2 remain to be discovered. Other antiviral factors that are not induced by IFN may also inhibit SARS-CoV-2: for instance, the RNA helicase DDX42 restricts several RNA viruses, including SARS-CoV-2 (23). While several whole-genome CRISPR/Cas9 screens identified host factors required for SARS-CoV-2 replication (24–29), none focused on antiviral genes.

Here, we performed a CRISPR/Cas9 screen designed to identify restriction factors for SARS-CoV-2, assessing the ability of 1905 ISGs to modulate SARS-CoV-2 replication in human epithelial lung cells. We report that the Death domain-associated protein 6 (DAXX), a scaffold protein residing in PML nuclear bodies (30) and restricting DNA viruses (31) as well as retroviruses (32,33), is a potent inhibitor of SARS-CoV-2 replication. SARS-CoV-2 restriction by DAXX is largely independent of the action of IFN, and unlike most of its other known activities, of the SUMOylation pathway. Within hours of infection, DAXX re-localizes to sites of viral replication in the cytoplasm, likely targeting viral transcription. We show that the SARS-CoV-2 papain-like protease (PLpro) induces the proteasomal degradation of DAXX, demonstrating that SARS-CoV-2 developed a mechanism to evade, at least partially, the restriction imposed by DAXX.

## Results

### A restriction factor-focused CRISPR/Cas9 screen identifies genes potentially involved in SARS-CoV-2 inhibition

To identify restriction factors limiting SARS-CoV-2 replication, we generated a pool of A549-ACE2 cells knocked-out (KO) for 1905 potential ISGs, using the sgRNA library we previously developed to screen HIV-1 restriction factors (34). This library includes more ISGs than most published libraries, as the inclusion criteria was less stringent (fold-change in gene expression in THP1 cells, primary CD4+ T cells or PBMCs ≥ 2). Therefore, some genes present in the library may not be ISGs *per se* in A549 cells. Transduced cells were selected by puromycin treatment, treated with IFNα and infected with SARS-CoV-2. Infected cells were immuno-labelled with a spike (S)-specific antibody and analyzed by flow cytometry. As expected (11,12), IFNα inhibited infection by 7-fold (**Fig. S1**). Infected cells were sorted based on S expression (**Fig. 1a**), and DNA was extracted from infected and non-infected control cells. Integrated sgRNA sequences in each cell fraction were amplified by PCR and sequenced by Next Generation Sequencing (NGS). Statistical analyses using the MAGeCK package (35) led to the identification of sgRNAs significantly enriched or depleted in infected cells representing antiviral and proviral factors, respectively (**Fig. 1b**). Although our screen was not designed to explicitly study proviral factors, we did successfully identify the well-described SARS-CoV-2 co-factor cathepsin L (CTSL) (36), validating our approach. USP18, a negative regulator of the IFN signaling pathway (37), and ISG15, which favors Hepatitis C Virus replication (38), were also identified as proviral ISGs. Core IFN pathway genes such as the IFN receptor (IFNAR1), STAT1, and STAT2, were detected as antiviral factors, further validating our screening strategy. LY6E, a previously described inhibitor of SARS-CoV-2 entry (15,16), was also a significant hit. Moreover, our screen identified APOL6, IFI6, DAXX and HERC5, genes that are known to encode proteins with antiviral activity against other viruses (39–42), but had not previously been studied in the context of SARS-CoV-2 infection. For all these genes except APOL6, individual sgRNAs were consistently enriched (for antiviral factors) or depleted (for proviral factors) in the sorted population of infected cells, while non-targeting sgRNAs were not (**Fig. 1c**).

**Figure 1:**
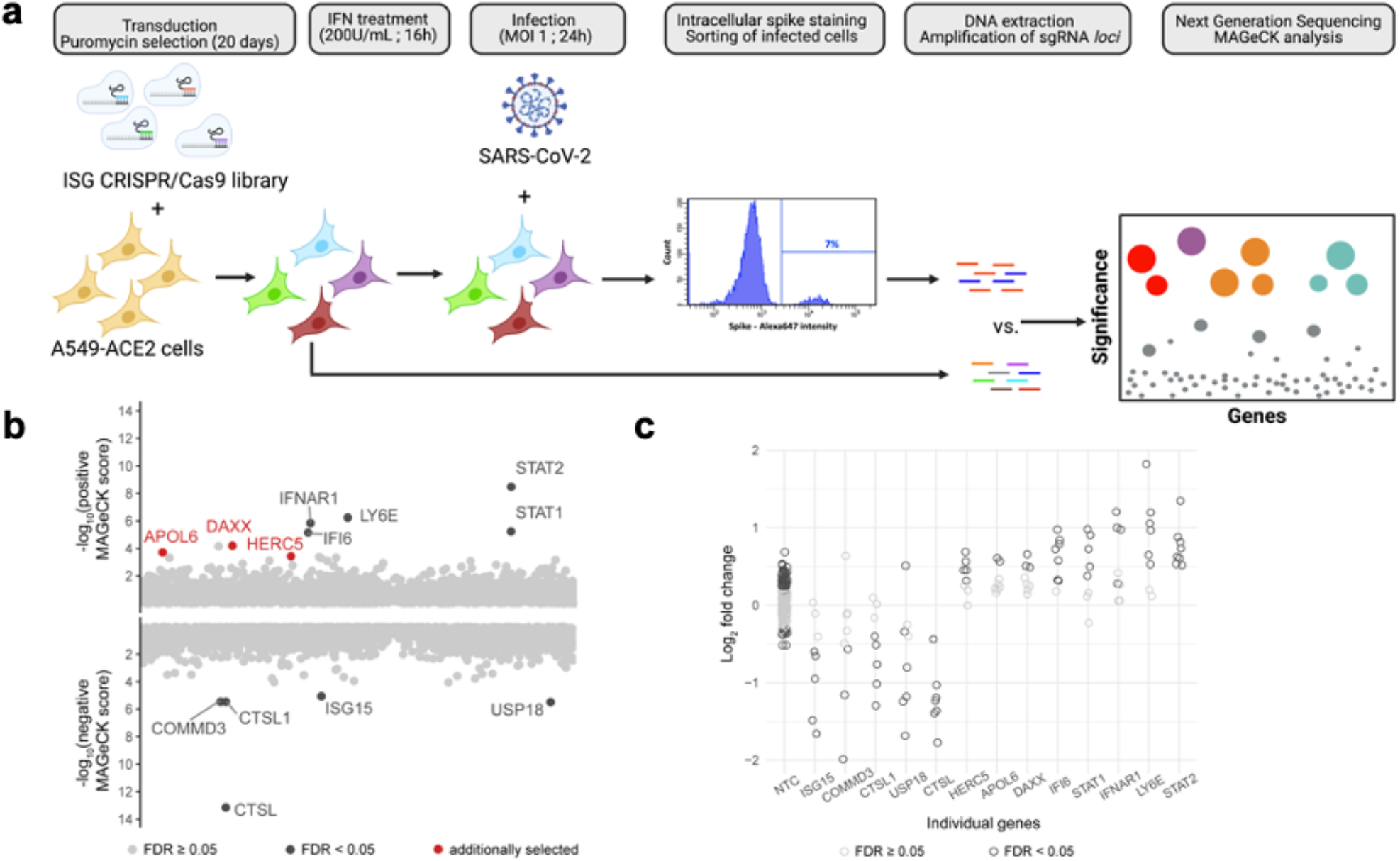
ISG-focused CRISPR/Cas9 screening approach to identify restriction factors for SARS-CoV-2. **a: CRISPR/Cas9 screen outline**. A549-ACE2 cells were transduced with lentivectors encoding the ISG CRISPR/Cas9 library and selected by puromycin treatment for 20 days. Library cells were then pre-treated with 200 U/mL of IFNα for 16 hours, and infection with SARS-CoV-2 at MOI 1. At 24h p.i., infected cells were fixed with formalin treatment, permeabilized by saponin treatment and stained with a monoclonal anti-spike antibody. After secondary staining, infected cells were sorted and harvested. Non-infected, non-IFNα treated cells were harvested as a control. DNA was extracted from both cellular fractions and sgRNA *loci* amplification was carried out by PCR. Following NGS, bio-informatic analysis using the MAGeCK package was conducted. **b: Screen results**. By taking into account the enrichment ratios of each of the 8 different sgRNAs for every gene, the MAGeCK analysis provides a positive score for KO enriched in infected cells (*i.e.* restriction factor, represented in the top fraction of the graph) and a negative score for KO depleted in infected cells (*i.e.* proviral factors, represented in the bottom portion of the graph). Gene with an FDR < 0.05 are represented in black. 3 genes with a FDR > 0.05, but with a p-value < 0.005 were additionally selected and are represented in red. **c: Individual sgRNA enrichment.** For the indicated genes, the enrichment ratio of the 8 sgRNAs present in the library was calculated as the MAGeCK normalized read counts in infected cells divided by those in the original pool of cells and is represented in log2 fold change. As a control, the enrichment ratios of the 200 non-targeting control (NTCs) is also represented.

### LY6E and DAXX display antiviral activity against SARS-CoV-2

To validate the ability of the identified hits to modulate SARS-CoV-2 replication in human cells, we generated pools of A549-ACE2 knocked-out (KO) cells for different genes of interest by electroporating a mix of 3 sgRNA/Cas9 ribonucleoprotein (RNP) complexes per gene target. Levels of gene editing were above 80% in all of the A549-ACE2 KO cell lines, as assessed by sequencing of the edited *loci* (**Table 1**). As controls, we used cells KO for IFNAR1, for the proviral factor CTSL or for the antiviral factor LY6E, as well as cells electroporated with non-targeting (NTC) sgRNAs/Cas9 RNPs. These different cell lines were then treated with IFNα and infected with SARS-CoV-2. Viral replication was assessed by measuring the levels of viral RNA in the supernatant of infected cells using RT-qPCR (**Fig. 2a**). In parallel, we titrated the levels of infectious viral particles released into the supernatant of infected cells (**Fig. 2b**). As expected, infection was significantly reduced in CTSL KO cells, confirming the proviral effect of this gene (36). Among the selected antiviral candidate genes, only 2 had a significant impact on SARS-CoV-2 replication: LY6E (as expected), and to an even greater degree, DAXX. Both genes restricted replication in absence of IFNα, an effect which was detectable at the level of viral RNA (8-fold and 42-fold reduction of infection, respectively, **Fig. 2a**) and of infectious virus (15-fold and 62-fold reduction, **Fig. 2b**). Based on available single-cell RNAseq datasets (43), DAXX is expected to be expressed in cell types physiologically relevant for SARS-CoV-2 infection such as lung epithelial cells and macrophages (**Fig. S2**).

**Table 1:** Gene editing efficiency. The frequency of editing was determined using Sanger sequencing and ICE analysis. Values are represented as mean ± SD (n=3).

| Gene | % of alleles edited |
| --- | --- |
| LY6E | 96 ± 1.73 |
| DAXX | 79,67 ± 2.52 |
| APOL6 | 99 ± 0 |
| HERC5 | 97 ± 0 |
| CTSL | 91 ± 1 |
| IFI6 | 88,33 ± 0.58 |
| IFNAR1 | 76,67 ± 3.21 |

**Figure 2:**
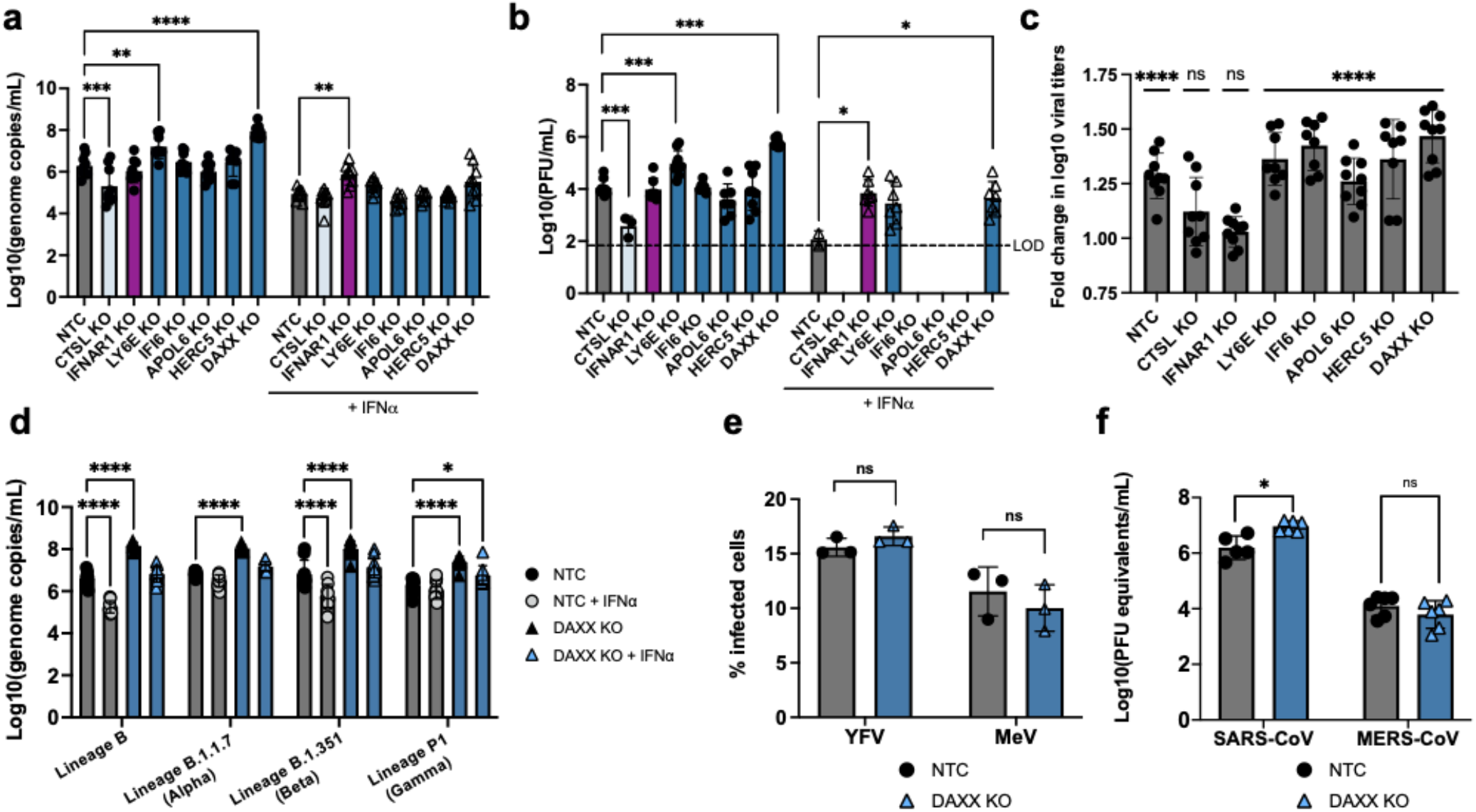
DAXX is a restriction factor for SARS-CoV-2. **a-c: Antiviral activity of ISGs against SARS-CoV-2.** A549-ACE2 knocked-out for the indicated genes were generated using a multi-guide approach, leading to pools of KO cells with a high frequency of indels. KO cells were pre-treated with 0 (circles) or 200 (triangles) U/mL of IFNα 24h prior to triplicate infection with SARS-CoV-2 (MOI 0.1). Supernatants were harvested at 72h p.i. The mean of three independent experiments, with infections carried out in triplicate, is shown. **a:** For the titration of RNA levels, supernatants were heat inactivated prior to quantification by qRT-PCR. Genome copies/mL were calculated by performing serial dilutions of a synthetic RNA with a known concentration. Statistics: 2-way ANOVA using Dunnett’s test, * = p-value < 0.05, ** = p-value < 0.01, *** = p-value < 0.001, **** = p-value < 0.0001. **b:** For the titration of infectious virus levels by plaque assay, supernatants were serially diluted and used to infect VeroE6 cells. Plaques formed after 3 days of infection were quantified using crystal violet coloration. Statistics: Dunnett’s test on a linear model, * p-value < 0.05, ** p-value < 0.01, *** p-value < 0.001. **c:** For each of the indicated KO, the data shown in **a.** is represented as fold change in log10 titers (*i.e.* the triplicate log10 titers of the non-treated condition divided by the mean of the triplicate log10 titers IFNα-treated condition, n=3). Statistics: 2-way ANOVA using Sidak’s test, ns = p-value > 0.05, **** = p-value < 0.001. **d-f: Antiviral activity of DAXX against SARS-CoV-2 variants and other viruses. d:** A549-ACE2 WT or DAXX KO cells were infected in triplicates at an MOI of 0.1 with the following SARS-CoV-2 strains: Lineage B (original strain); Lineage B.1.1.7. (Alpha variant); Lineage B.1.35.1 (Beta variant); Lineage P1 (Gamma variant). Supernatants were harvested at 72h p.i. Supernatants were heat inactivated prior to quantification by qRT-PCR. Genome copies/mL were calculated by performing serial dilutions of a synthetic RNA with a known concentration. The mean of three independent experiments, with infections carried out in triplicate, is shown. **e:** A549-ACE2 WT or DAXX KO cells were infected in triplicates with Yellow Fever Virus (YFV, Asibi strain, MOI of 0.3) or with Measles Virus (MeV, Schwarz strain expressing GFP, MOI of 0.2). At 24h p.i., the percentages of cells positive for viral protein E (YFV) or GFP (MeV) was assessed by flow cytometry. The mean of 3 independent experiments is represented. **f:** WT or DAXX KO cells were infected in triplicates at an MOI of 0.1 with SARS-CoV or MERS-CoV. Supernatants were harvested at 72h p.i. Supernatants were heat inactivated prior to quantification by qRT-PCR. Serial dilutions of a stock of known infectious titer was used as a standard. The mean of 2 independent experiments is represented. Statistics: 2-way ANOVA using Dunnett’s test, * = p-value < 0.05, *** = p-value < 0.001, **** = p-value < 0.0001

In IFNα-treated cells, DAXX and LY6E KO led to a modest, but significant rescue of viral replication, which was particularly visible when measuring the levels of infectious virus by plaque assay titration (**Fig. 2b**), while the antiviral effect of IFNα treatment was completely abrogated in IFNAR1 KO cells, as expected (**Fig. 2c**). However, IFNα still had robust antiviral effect on SARS-CoV-2 replication in both DAXX KO and LY6E KO cells (**Fig. 2c**). While DAXX and LY6E contribute to the IFN-mediated restriction, this suggests that there are likely other ISGs contributing to this effect. DAXX is sometimes referred to as an ISG, and was originally included in our ISG library, although its expression is only weakly induced by IFN in some human cell types (32,44). Consistent with this, we found little to no increase in DAXX expression in IFNα-treated A549-ACE2 cells (**Fig. S3**). In addition, we tested the antiviral effect of DAXX on several SARS-CoV-2 variants that have been suggested to be partially resistant to the antiviral effect of IFN in A549-ACE2 cells (45). Our results confirmed that Lineage B.1.1.7. (Alpha) and Lineage P1 (Gamma) SARS-CoV-2 variants were indeed less sensitive to IFN (**Fig. 2d**). DAXX, however, restricted all variants to a similar level than the original Lineage B strain of SARS-CoV-2 (**Fig. 2d**), suggesting that while some variants may have evolved towards IFN-resistance, they are still efficiently restricted by DAXX. To determine whether DAXX is specific to SARS-CoV-2 or also inhibits other RNA viruses, including coronaviruses, we infected A549-ACE2 WT and DAXX KO cells with SARS-CoV, MERS-CoV, and 2 RNA viruses belonging to unrelated viral families: Yellow Fever Virus (YFV) and Measles Virus (MeV), which are positive and negative strand RNA viruses, respectively. Our results show that DAXX restricts SARS-CoV, but has no effect on the replication of YFV, MeV or MERS-CoV (**Fig. 2e-f**). Thus, our data suggests that DAXX restriction may exhibit some level of specificity.

### DAXX targets SARS-CoV-2 transcription

Next, we investigated whether DAXX targets early steps of the SARS-CoV-2 viral life cycle such as viral entry or transcription. The intracellular levels of two viral transcripts were assessed at different time post-infection in A549-ACE2 WT or DAXX KO cells (**Fig. 3**). At early time points (from 2h to 6h p.i.), the levels of viral RNA were similar in WT and DAXX KO cells, suggesting that comparable amounts of SARS-CoV-2 virions were entering cells. The levels of viral transcripts significantly increased starting at 8h p.i., representing the initiation of viral transcription. The levels of the 5’ UTR viral transcript (**Fig. 3a)** were 6.4-fold higher at 8h; 4.1-fold higher at 10h; and 8-fold higher at 24h post-infection in DAXX KO cells compared to WT cells. The levels of RdRp transcripts were less affected by the absence of DAXX than 5’UTR transcripts (**Fig. 3b**) with levels of viral transcripts 1.7-fold and 3.5-fold higher in DAXX KO cells compared to WT cells at 10h and 24h pos-infection, respectively. These results suggest that DAXX acts early during the SARS-CoV-2 replication cycle, likely targeting the step of viral transcription.

**Figure 3:**
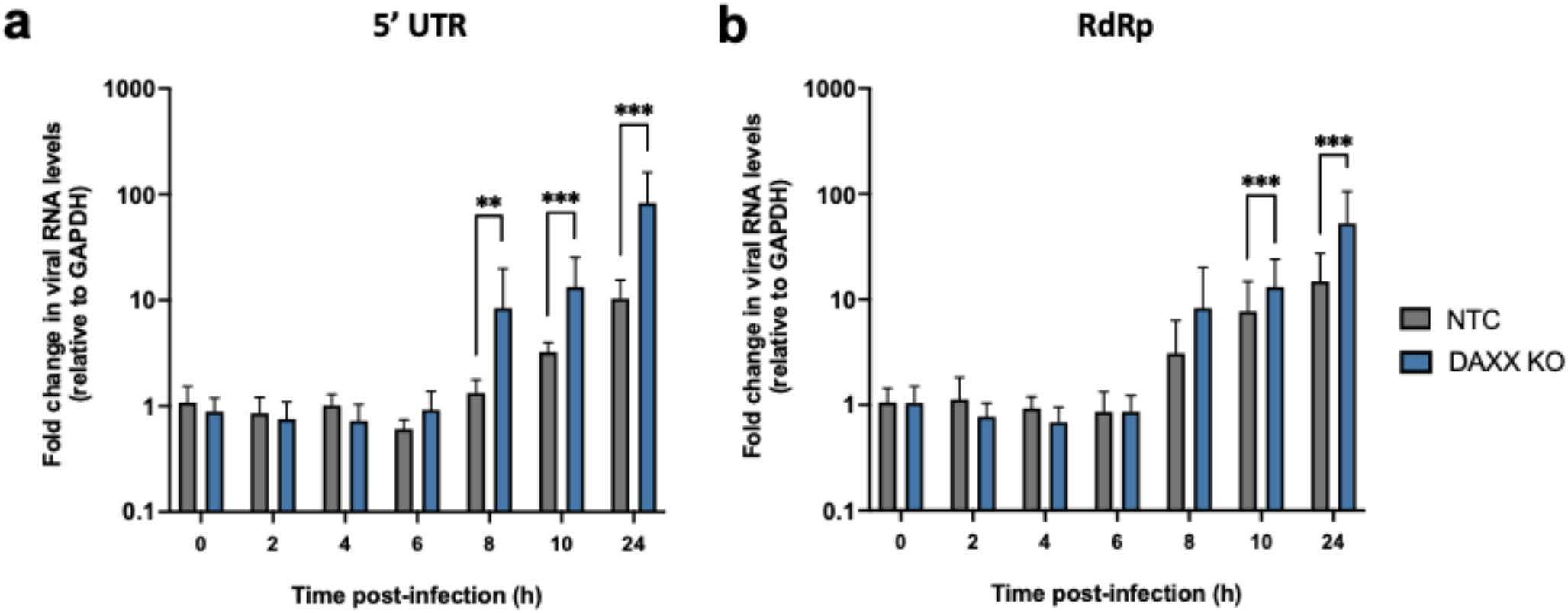
DAXX restricts SARS-CoV-2 transcription. A549-ACE2 WT or DAXX KO were infected at MOI 1 in triplicates. Cell monolayers were harvested at the indicated time points, and total RNA was extracted. The levels of viral RNA (**a:** 5’ UTR; **b:** RdRp) were determined by qRT-PCR and normalized against GAPDH levels. The mean of 3 independent experiments is represented. Statistics: Dunnett’s test on a linear model, * p-value < 0.05, ** p-value < 0.01, *** p-value < 0.001.

### DAXX restriction is SUMO-independent

DAXX is a small scaffold protein that acts by recruiting other SUMOylated proteins in nuclear bodies through its C-terminal SUMO-Interacting Motif (SIM) domain (46). The recruitment of these factors is required for the effect of DAXX on various cellular processes such as transcription and apoptosis, and on its antiviral activities (32,47–49). DAXX can also be SUMOylated itself (50), which may be important for some of its functions. To investigate the role of SUMOylation in DAXX-mediated SARS-CoV-2 restriction, we used overexpression assays to compare the antiviral activity of DAXX WT with two previously described DAXX mutants (51). First, we used a version of DAXX in which 15 lysine residues have been mutated to arginine (DAXX 15KR), which is unable to be SUMOylated; and second, a truncated version of DAXX that is missing its C-terminal SIM domain (DAXXΔSIM) (48) and is unable to interact with its SUMOylated partners. A549-ACE2 were refractory to SARS-CoV-2 infection upon transfection with any plasmid, precluding us from using this cell line. Instead, we transfected 293T-ACE2 cells, another SARS-CoV-2 permissive cell line (18).

We examined the effect of DAXX WT overexpression on the replication of SARS-CoV-2-mNeonGreen (52) by microscopy. DAXX overexpression starkly reduced the number of infected cells (**Fig. 4a-b**), revealing that DAXX-mediated restriction is not specific to A549-ACE2 cells. Using double staining for HA-tagged DAXX and SARS-CoV-2, we found that most of the DAXX-transfected cells were negative for infection, and conversely, that most of the infected cells did not express transfected DAXX (**Fig. 4c**), indicating that DAXX imposes a major block to SARS-CoV-2 infection.

**Figure 4:**
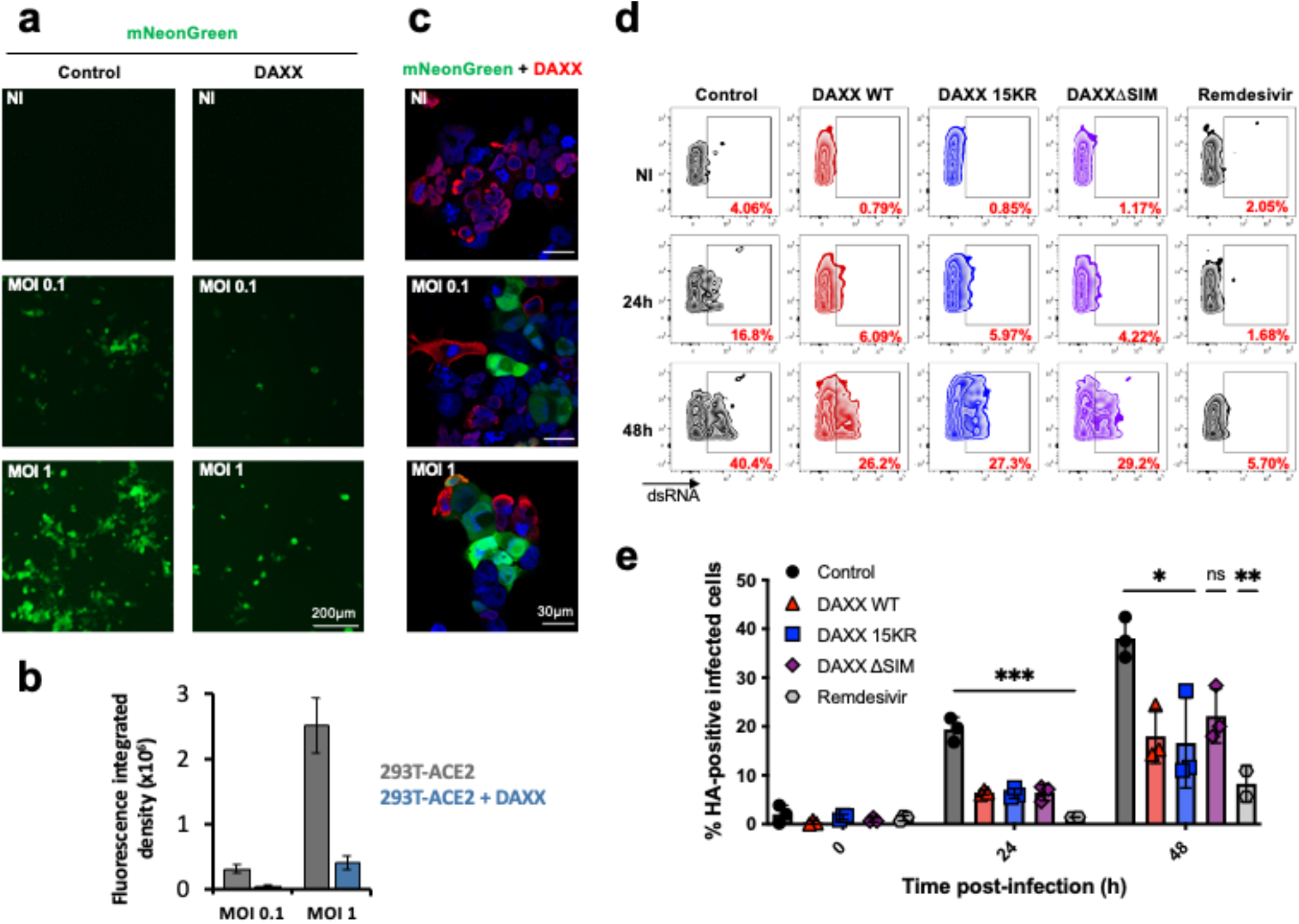
DAXX restriction of SARS-CoV-2 is SUMOylation independent. **a-c: DAXX overexpression restricts SARS-CoV-2.** 293T-ACE2 cells were transfected with DAXX WT. 24h after transfection, cells were infected with the mNeonGreen fluorescent reporter SARS-CoV-2 at the indicated MOI. Cells were either visualized with an EVOS fluorescence microscope (**a-b**) or stained with an HA-antibody detecting DAXX and imaged by confocal microscopy (**c**). Scale bars correspond to 200 μm (**a)** and 30 μm (**c**). Images shown in (**a**) were quantified using ImageJ software (**b**). Data show the mean +/- SD of Fluorescence integrated densities. The analysis was performed on around 200 cells from 3 different fields. **d-e: DAXX mutants are still able to restrict SARS-CoV-2**. 293T-ACE2 cells were transfected with HA-DAXX WT; HA-DAXX 15KR; HA-DAXXΔSIM; or with HA-NBR1 as negative control plasmid. 24h after transfection, cells were infected with SARS-CoV-2 at an MOI of 0.1. When indicated, cells were treated with remdesivir at the time of infection. After 24h or 48h, infected cells were double-stained recognizing dsRNAs (to read out infection) and HA (to read out transfection efficiency) and acquired by flow cytometry. The percentage of infected cells among HA-positive (transfected) cells for one representative experiment is shown in **d**, for the mean of 3 independent experiments in **e**. Statistics: one-way ANOVA using Dunnett’s test, Holm corrected, ns = p-value > 0.05, * = p-value < 0.05, ** = p-value < 0.01, *** = p-value < 0.001.

In order to quantify the antiviral effect of overexpressed DAXX WT and mutants, we assessed the number of cells positive for the S protein (among transfected cells) by flow cytometry and the abundance of viral transcripts by qRT-PCR. Western blot (**Fig. S4a**) and flow cytometry (**Fig. S4b**) analyses showed that DAXX WT and mutants were expressed at similar levels, with a transfection efficiency of around 40 to 50% for all three constructs. DAXX WT, 15KR and ΔSIM all efficiently restricted SARS-CoV-2 replication. Indeed, at 24 hours p.i., the proportion of infected cells (among HA-positive cells) was reduced by 2 to 3-fold as compared to control transfected cells for all 3 constructs (**Fig. 4d**). This effect was less pronounced but still significant at 48 hours p.i. (**Fig. 4e**). Moreover, DAXX overexpression led to a significant reduction of the levels of two different viral transcripts (**Fig. S5**), in line with our earlier results showing that DAXX targets viral transcription (**Fig. 3a-b**). Together, these results show that DAXX overexpression restricts SARS-CoV-2 replication in a SUMOylation-independent mechanism.

### SARS-CoV-2 infection triggers DAXX re-localization

DAXX mostly localizes in nuclear bodies (30), whereas SARS-CoV-2 replication occurs in the cytoplasm. We reasoned that DAXX localization may be altered during the course of infection in order for the restriction factor to exert its antiviral effect. To test this hypothesis, we infected 293T-ACE2 cells with SARS-CoV-2 and used high-resolution confocal microscopy to study the localization of endogenous DAXX (**Fig. 5**). As expected (30), DAXX mostly localizes in the nuclei of non-infected cells, forming discrete *foci*. At 6h post-infection, DAXX re-localizes to the cytoplasm, although nuclear *foci* can still be detected. At 24h post-infection, DAXX is completely depleted from nuclear bodies, and is found almost exclusively in the cytoplasm of infected cells, in close association with in close association with dsRNAs, likely representing SARS-CoV-2 viral dsRNAs. These results suggest that early events following SARS-CoV-2 infection trigger the re-localization of DAXX from the nucleus to the cytoplasm.

**Figure 5:**
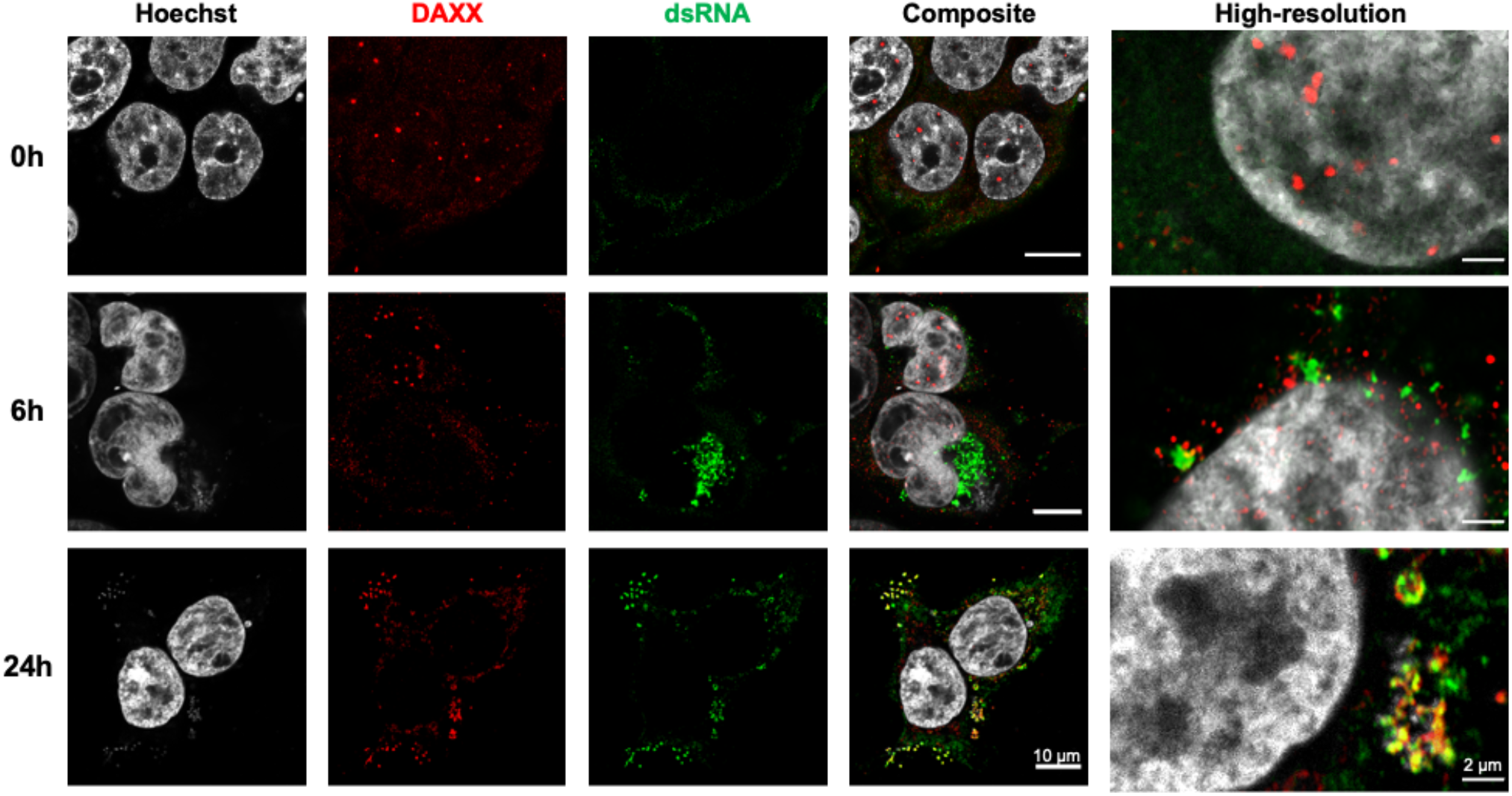
SARS-CoV-2 infection induces DAXX cytoplasmic re-localization to sites of viral replication. 293T-ACE2 cells were infected with SARS-CoV-2 at the indicated MOI 1. 24h post-infection, cells were labelled with Hoescht and with antibodies against dsRNA (detecting viral RNA, in green) and HA (detecting DAXX, in red). When indicated, the high-resolution Airyscan mode was used. Scale bars correspond to 10 μm for confocal images, and 2 μm for the high-resolution images.

### SARS-CoV-2 PLpro induces proteasomal degradation of DAXX

Next, we asked whether this relocalization of DAXX following infection destabilizes the protein. Western blot analysis revealed that SARS-CoV-2 infection induces a marked decrease of total DAXX expression in infected cells (**Fig. 6a**). In contrast, SARS-CoV-2 infection had no effect on DAXX mRNA levels (**Fig. S6**). Importantly, the decrease in DAXX protein levels is likely not attributed to a global host cell shut down, as the levels of Lamin B, HSP90, Actin, GAPDH, Tubulin, TRIM22 and RIG-I were unchanged following infection (**Fig. 6a**). These results suggest that DAXX may be actively and specifically targeted by SARS-CoV-2 for degradation during the course of infection. SARS-CoV-2 papain-like protease (PLpro) is a possible candidate for this activity, as it cleaves other cellular proteins such as ISG15 (53,54), and ULK1 (55). It was also shown that foot-and-mouth disease virus (FDMV) PLpro degrades DAXX (56). Thus, we treated cells with different inhibitors: GRL0617, an inhibitor of SARS-CoV-2 PLpro (54); MG132, a well-described proteasome inhibitor; or Masitinib, an inhibitor of SARS-CoV-2 3CL protease (57). These inhibitors had minimal effects on cell viability at the selected concentrations (**Fig**. **S7**). Strikingly, GRL0617 treatment partially restored DAXX expression (**Fig. 6b**), especially at the highest concentration. Similarly, MG132 also prevented DAXX degradation in SARS-CoV-2 infected cells. In contrast, Masitinib treatment had no effect on DAXX levels. These results suggest that PLpro, but not 3CL, targets DAXX for proteasomal degradation. Consistently, GRL0617 treatment also restored DAXX subcellular localization to nuclear bodies (**Fig. 6c**). As expected, GRL0617 treatment also inhibited the production of SARS-CoV-2 proteins such as spike (**Fig. 6b**), and may thus have an indirect effect on DAXX by inhibiting SARS-CoV-2 replication itself. However, the fact that Masitinib also inhibits spike production but does not restore DAXX expression suggested that DAXX degradation is not an unspecific consequence of viral replication but rather a specific activity of PLpro. To investigate the potential direct contribution of PLpro to DAXX degradation, we assessed the impact of overexpressing individual SARS-CoV-2 proteins in 293T-ACE2 cells on DAXX levels. We included in the analysis mCherry-tagged SARS-CoV-2 Non-structural proteins (Nsp) (58), which are not expressed from a lentiviral vector that may be targeted by DAXX antiviral activity (33). This included Nsp3 (which encodes PLro), Nsp4, Nsp6, Nsp7, Nsp10, Nsp13 and Nsp14. All proteins were expressed at similar levels (**Fig. S8a**). Only the overexpression of Nsp3 led to DAXX degradation (**Fig. 6d**). This effect was dose-dependent (**Fig. 6e and Fig. S8b**), and was abrogated when cells were treated with GRL0617 (**Fig. 6f**). Taken together, these results strongly indicate that PLpro directly induces the proteasomal degradation of DAXX.

**Figure 6:**
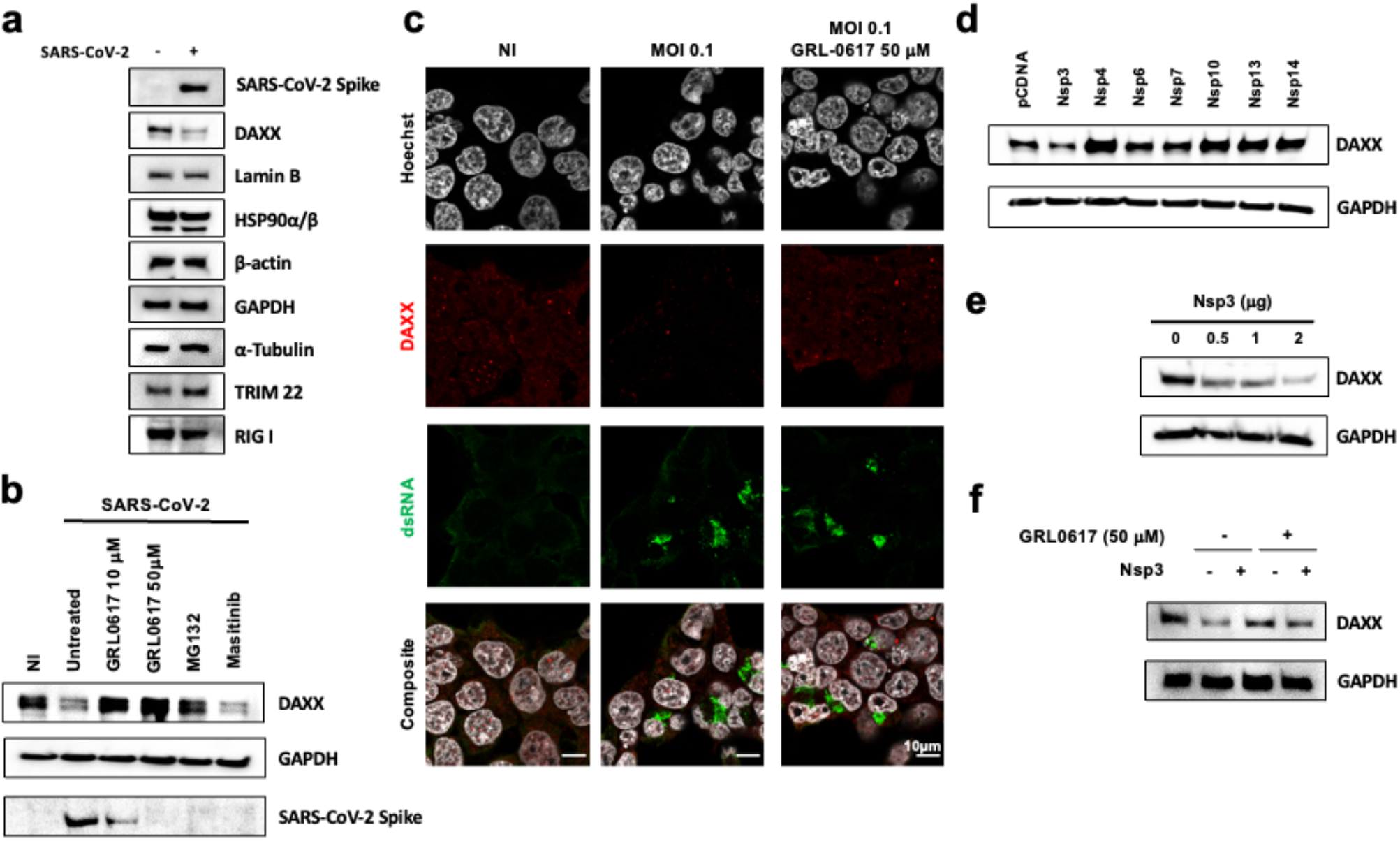
SARS-CoV-2 PLpro induces the proteasomal degradation of DAXX. **a: DAXX degradation after infection.** 293T-ACE2 cells were infected with SARS-CoV-2 at MOI 0.1. After 24h, cells were harvested and levels of DAXX, Lamin B, HSP90, Actin, GAPDH, Tubulin, TRIM22, RIG-I and of the viral protein spike were analyzed by Western Blot. **b: GRL0617 and MG132 treatment restores DAXX expression.** 293T-ACE2 cells were infected with SARS-CoV-2 at MOI 0.1. When indicated, cells were pretreated 2h before infection with GRL0617 (at the indicated concentrations), or with MG132 (10 μM), a proteasome inhibitor, or Masitinib (10 μM) a 3CL inhibitor. After 24h, cells were harvested and levels of DAXX, GAPDH and of the viral protein spike were analyzed by Western Blot. **c: GRL0617 treatment restores DAXX localization.** 293T-ACE2 cells were infected with SARS-CoV-2 at MOI 0.1. 24h post-infection, cells were labelled with Hoescht and with antibodies against dsRNA (detecting viral RNA, in green) and HA (detecting DAXX, in red). When indicated, cells were treated with 50 μM of GRL0617 at the time of infection. Scale bars correspond to 10 μm. **d-f: Nsp3 induces DAXX degradation. D:** 293T-ACE2 cells were transfected with 1 μg of the indicated viral proteins. After 24h, the levels of DAXX and GAPDH were analyzed by Western Blot. **E:** 293T-ACE2 cells were transfected with the indicated amounts of Nsp3. After 24h, the levels of DAXX and GAPDH were analyzed by Western Blot. **f:** 293T-ACE2 cells were transfected with 1 μg of Nsp3. 6 hours post transfection, cells were also, when indicated, treated with 50 μM of GRL0617. 24h after transfection, the levels of DAXX and GAPDH were analyzed by Western Blot.

## Discussion

### Comparison with other screens

The whole-genome CRISPR/Cas9 screens conducted to date on SARS-CoV-2 infected cells mostly identified host factors necessary for viral replication (24–29) and did not focus on antiviral genes, as did our screen. Three overexpression screens, however, identified ISGs with antiviral activity against SARS-CoV-2 (16,22,21). In the first one, Pfaender *et al.* screened 386 ISGs for their antiviral activity against the endemic human coronavirus 229E, and identified LY6E as a restriction factor inhibiting both 229E and SARS-CoV-2. Our screen also identified LY6E as a top hit (**Fig.1**), further validating the findings of both studies. Four additional genes had significant p-values in both Pfaender *et al.* and our work: IFI6, HERC5, OAS2 and SPSB1 (**Table S5-S6**). We showed that knocking-out LY6E and DAXX only partially rescued SARS-CoV-2 replication in IFN-treated cells (**Fig. 2)**, suggesting that other IFN effectors active against SARS-CoV-2 remain to be identified. For instance, other ISGs, such as IFITMs, inhibit SARS-CoV-2 viral entry (17–19). In the second screen, Martin Sancho *et al.* tested 399 ISGs against SARS-CoV-2. Among the 65 antiviral ISGs identified, they focused on BST-2/Tetherin, which targets viral budding. BST-2/Tetherin was not a significant hit in our screen (**Table S5-6**). This discrepancy is likely due to the fact that our screen relies on the sorting of S-positive cells, and is therefore unable to detect factors restricting the late stages of the viral replication cycle. The most recent overexpression screen assessed the contribution of 539 human and 444 macaque ISGs in SARS-CoV-2 restriction, and further characterized the role of OAS1 in sensing SARS-CoV-2 and restricting its replication through RNAseL. While we did not identify OAS1 or RNAseL in our screen (**Table S5-6**), we did identify hits in common with this screen, including IFI6 and OAS2 (that were also identified by Pfaender *et al.*). Of note, DAXX was absent from the ISG libraries used by these overexpression screens, which explains why it was not previously identified as an antiviral gene for SARS-CoV-2. Our sgRNA library, by including 1905 genes, targeted a wider set of ISGs and “ISG-like” genes, including genes like DAXX that are not (or only weakly) induced by IFN in some cell types (32,44). Interestingly, IFN has a stronger effect on DAXX expression levels in cells from other mammals such as bats (59). Future studies may investigate whether DAXX orthologs of different species are also able to restrict SARS-CoV-2 and whether DAXX participates in IFN-mediated viral restriction in these species.

### DAXX is a restriction factor for SARS-CoV-2

We identify DAXX as a potent antiviral factor restricting the replication of SARS-CoV-2, acting independently of IFN and likely targeting an early step of the viral life cycle such as transcription (**Fig. 3**). DAXX fulfills all of the criteria defining a *bona fide* SARS-CoV-2 restriction factor: knocking-out endogenous DAXX leads to enhanced viral replication (**Fig. 2**), while over-expression of DAXX restricts infection (**Fig. 4**). DAXX co-localizes with viral dsRNAs (**Fig. 5**) and SARS-CoV-2 antagonizes DAXX to some extent, as evidenced by the proteasomal degradation of DAXX induced by PLpro (**Fig. 6**). Although DAXX expression is not upregulated by IFNα in A549 cells (**Fig. S3**), basal levels of expression are sufficient for its antiviral activity, as has been shown for other potent restriction factors. Publicly available single-cell RNAseq analyses (**Fig. S2**) indicated that DAXX is expressed in cell types targeted by the virus *in vivo*, such as lung epithelial cells and macrophages. Interestingly, DAXX exhibited some degree of specificity in its antiviral activity, as unrelated viruses such as YFV and MeV, as well as the closely related MERS-CoV were not sensitive to its action, in contrast to SARS-CoV and SARS-CoV-2 (**Fig. 2**). Future work will determine which viral determinants are responsible for this specific antiviral activity of DAXX.

### Mechanism of DAXX-mediated restriction

DAXX is mostly known for its antiviral activity against DNA viruses replicating in the nucleus, such as adenovirus 5 (AdV5) (60) and human papillomavirus (HPV) (61). Most of these viruses antagonize PML and/or DAXX, which interacts with PML in nuclear bodies (30). We show here that DAXX is also able to restrict SARS-CoV-2, a positive sense RNA virus that replicates in the cytoplasm. Recent studies have shown that DAXX inhibits the reverse transcription of HIV-1 in the cytoplasm (32,33). Within hours of infection, DAXX subcellular localization was altered, with DAXX accumulating in the cytoplasm and colocalizing with incoming HIV-1 capsids (33). Here, we observed a similar phenomenon, with a rapid re-localization of DAXX from the nucleus to cytoplasmic viral replication sites (**Fig. 5**), where it likely exerts its antiviral effect. Early events in the replication cycle of both HIV-1 and SARS-CoV-2, such as viral fusion or virus-induced stress, may thus trigger DAXX re-localization to the cytoplasm. DAXX seems to inhibit SARS-CoV-2 by a distinct mechanism: whereas the recruitment of interaction partners through the SIM-domain is required for the effect of DAXX on HIV-1 reverse transcription (32), it was not the case in the context of SARS-CoV-2 restriction. This result was unexpected, since DAXX has no enzymatic activity and rather acts as a scaffold protein recruiting SUMOylated partners through its SIM domain (51). Some DAXX functions, such as interaction with the chromatin remodeler ATRX (30) or its recently described role as a chaperone protein (62) are, however, SIM-independent. Future work should determine which DAXX domains and residues are required for its antiviral activity.

### Antagonism of DAXX by SARS-CoV-2

Our results suggest that SARS-CoV-2 developed a mechanism to antagonize DAXX restriction, with PLpro inducing its degradation to the proteasome (**Fig. 6**). his antagonism, however, is only partial, since knocking-out DAXX still enhances SARS-CoV-2 replication (**Fig. 2**). Another possibility is that DAXX, by acting early in the viral life cycle (i.e. as soon as 8 hours p.i., **Fig. 3**) may exert its antiviral effect before PLpro is able to complete its degradation. Proteins expressed by other viruses are also able to degrade DAXX: for instance, the AdV5 viral factor E1B-55K targets DAXX for proteasomal degradation (60), and FDMV PLpro cleaves DAXX (56). We showed in **Fig. 2** that SARS-CoV, but not MERS-CoV, is sensitive to DAXX. Thus, it will be interesting to test whether PLpro from these different coronaviruses differ in their ability to degrade DAXX, and whether this has an impact on their sensitivity to DAXX restriction. Future research may also establish whether PLpro induces the degradation of DAXX through direct cleavage, or whether it acts in a more indirect way, such as cleaving or recruiting cellular co-factors. Such investigations may be relevant for the development of PLpro inhibitors (63): indeed, in addition to directly blocking SARS-CoV-2 replication, PLpro inhibitors may also sensitize the virus to existing antiviral mechanisms such as DAXX restriction.

## Material & Methods

### Cells, viruses & plasmids

HEK 293T (ATCC #CRL-11268) were cultured in MEM (Gibco #11095080) complemented with 10% FBS (Gibco #A3160801) and 2 mM L-Glutamine (Gibco # 25030081). VeroE6 (ATCC #CRL-1586), A549 (ATCC #CCL-185) and HEK 293T, both overexpressing the ACE2 receptor (A549-ACE2 and HEK 293T-ACE2, respectively), were grown in DMEM (Gibco #31966021) supplemented with 10% FBS (Gibco #A3160801), and penicillin/streptomycin (100 U/mL and 100 μg/mL, Gibco # 15140122). Blasticidin (10 μg/mL, Sigma-Aldrich #SBR00022-10ML) was added for selection of A549-ACE2 and HEK 293T-ACE2. All cells were maintained at 37°C in a 5% CO_2_ atmosphere. Universal Type I Interferon Alpha (PBL Assay Science #11200-2) was diluted in sterile-filtered PBS 1% BSA according to the activity reported by the manufacturer. The strains BetaCoV/France/IDF0372/2020 (Lineage B); hCoV-19/France/IDF-IPP11324/2020 (Lineage B.1.1.7); and hCoV-19/France/PDL-IPP01065/2021 (Lineage B.1.351) were supplied by the National Reference Centre for Respiratory Viruses hosted by Institut Pasteur and headed by Pr. Sylvie van der Werf. The human samples from which the lineage B, B.1.1.7 and B.1.351 strains were isolated were provided by Dr. X. Lescure and Pr. Y. Yazdanpanah from the Bichat Hospital, Paris, France; Dr. Besson J., Bioliance Laboratory, saint-Herblain France; Dr. Vincent Foissaud, HIA Percy, Clamart, France, respectively. These strains were supplied through the European Virus Archive goes Global (Evag) platform, a project that has received funding from the European Union’s Horizon 2020 research and innovation programme under grant agreement #653316. The hCoV-19/Japan/TY7-501/2021 strain (Lineage P1) was kindly provided by Jessica Vanhomwegen (Cellule d’Intervention Biologique d’Urgence; Institut Pasteur). The mNeonGreen reporter SARS-CoV-2 was provided by Pei-Yong Shi (52). SARS-CoV FFM-1 strain (64) was kindly provided by H.W. Doerr (Institute of Medical Virology, Frankfurt University Medical School, Germany). The Middle East respiratory syndrome (MERS) Coronavirus, strain IP/COV/MERS/Hu/France/FRA2 (Genbank reference <u>KJ361503</u>) isolated from one of the French cases (65) was kindly provided by Jean-Claude Manuguerra (Cellule d’Intervention Biologique d’Urgence; Institut Pasteur). SARS-CoV-2 viral stocks were generated by infecting VeroE6 cells (MOI 0.01, harvesting at 3 dpi) using DMEM supplemented with 2% FBS and 1 μg/mL TPCK-trypsin (Sigma-Aldrich #1426-100MG). SARS-CoV and MERS-CoV viral stocks were generated by infecting VeroE6 cells (MOI 0.0001) using DMEM supplemented with 5% FCS and harvesting at 3 dpi or 6 dpi, respectively. The Yellow Fever Virus (YFV) Asibi strain was provided by the Biological Resource Center of the Institut Pasteur. The Measles Schwarz strain expressing GFP (MeV-GFP) was described previously (70). Both viral stocks were produced on Vero NK cells. The Human Interferon-Stimulated Gene CRISPR Knockout Library was a gift from Michael Emerman and is available on Addgene (Pooled Library #125753). The plentiCRISPRv.2 backbone was ordered through Addgene (Plasmid #52961). pMD2.G and psPAX2 were gifts from Didier Trono (Addgene #12259; #12260). pcDNA3.1 was purchased from Invitrogen. Plasmids constructs expressing WT and mutant HA-tagged DAXX constructs were kindly provided by Hsiu-Ming Shih (51). The plasmids encoding mCherry-tagged viral proteins were a gift from Bruno Antonny and ordered through Addgene: Nsp3-mCherry (#165131); Nsp4-mCherry (#165132); Nsp6-mCherry (#165133); Nsp7-mCherry (#165134); Nsp10-mCherry (#165135); Nsp13-mCherry (#165136); Nsp14-mCherry (#165137).

### Antibodies

For Western Blot, we used mouse anti-DAXX (diluted 1:1000, Abnova #7A11), rat anti-HA clone 3F10 (diluted 1:3000, Sigma #2158167001), mouse anti-GAPDH clone 6C5 (diluted 1:3000, Millipore #FCMAB252F), Goat anti-Lamin B clone M-20 (diluted 1:500, Santa Cruz sc-6217), mouse monoclonal HSP90α/β clone F-8 (diluted 1 :500, Santa Cruz sc-13119), mouse monoclonal β-actin clone AC-15 (1:3000 Sigma #A1978), mouse monoclonal α-Tubulin clone DMA1 (diluted 1:1000, Sigma #T9026), rabbit anti-TRIM22 (diluted 1 :1000, Proteintech #13744-1-AP) and mouse Monoclonal RIG-I clone Alme-1 (diluted 1: 1000, adipoGen #AG-20B-0009). To detect SARS-CoV-2 Spike protein, we used mouse anti-spike clone 1A9 (diluted 1:1000, GeneTex GTX632604). Secondary antibodies were goat anti-mouse and anti-rabbit HRP-conjugates (diluted 1:5000, ThermoFisher #31430 and #31460) and horse anti-goat HRP (diluted 1: 1000, Vector # PI-9500). For immunofluorescence, we used rabbit anti-DAXX (diluted 1:50, Proteintech #20489-1-AP) and mouse anti-dsRNA J2 (diluted 1:50, Scicons #10010200). Secondary antibodies were goat anti-rabbit AF555 and anti-mouse AF488 (diluted 1:1000, ThermoFisher #A-21428 and #A-28175). For flow sorting of infected cells, we used the anti-S2 H2 162 antibody (diluted 1:150), a kind gift from Dr. Hugo Mouquet, (Institut Pasteur, Paris, France). Secondary antibody was donkey anti-mouse AF647 (diluted 1:1000, Invitrogen #A31571). For FACS analysis, we used rat anti-HA clone 3F10 (diluted 1:100, Sigma #2158167001) and mouse anti-dsRNA J2 (diluted 1:500, Scicons #10010200). Secondary antibodies were goat anti-rat AF647 and anti-mouse AF488 (diluted 1:1000, ThermoFisher #A-21247 #A-28175). The pan-flavivirus anti-Env 4G2 antibody was a kind gift from Phillipe Desprès.

### Generation of CRISPR/Cas9 library cells

HEK 293T cells were transfected with the sgRNA library plasmid together with plasmids coding for Gag/Pol (R8.2) and for the VSVg envelope (pVSVg) using a ratio of 5:5:1 and calcium phosphate transfection. Supernatants were harvested at 36h and 48h, concentrated 80-fold by ultracentrifugation (22,000 g, 4°C for 1h) and pooled. To generate the ISG KO library cells, 36×10^6^ A549-ACE2 cells were seeded in 6 well plates (10^6^ cells per well) 24h before transduction. For each well, 100 μL of concentrated lentivector was diluted in 500 μL of serum-free DMEM, supplemented with 10 μg/mL of DEAE dextran (Sigma #D9885). After 48h, transduced cells were selected by puromycin treatment for 20 days (1 μg/mL; Sigma #P8833).

### CRISPR/Cas9 screen

4×10^7^ A549-ACE2 cells were treated with IFNα (200U/mL). 16h later, cells were infected at a MOI of 1 in serum-free media complemented with TPCK-trypsin and IFNα (200 U/mL). After 90 min, the viral inoculum was removed, and cells were maintained in DMEM containing 5% FBS and IFNα (200 U/mL). After 24h, cells were harvested and fixed for 15 min in Formalin 1%. Fixed cells were washed in cold FACS buffer containing PBS, 2% Bovine Serum Albumin (Sigma-Aldrich #A2153-100G), 2 mM EDTA (Invitrogen #15575-038) and 0.1% Saponin (Sigma-Aldrich #S7900-100G). Cells were incubated for 30 min at 4°C under rotation with primary antibody diluted in FACS buffer. Incubation with the secondary antibody was performed during 30 min at 4°C under rotation. Stained cells were resuspended in cold sorting buffer containing PBS, 2% FBS, 25 mM Hepes (Sigma-Aldrich #H0887-100ML) and 5 mM EDTA. Infected cells were sorted on a BD FACS Aria Fusion. Sorted and control (non-infected, not IFN-treated) cells were centrifugated (20 min, 2,000g) and resuspended in lysis buffer (NaCI 300 mM, SDS 0.1%, EDTA 10 mM, EGTA 20 mM, Tris 10 mM) supplemented with 1% Proteinase K (Qiagen #19133) and 1% RNAse A/T1 (ThermoFisher #EN0551) and incubated overnight at 65°C. Two consecutive phenol-chloroform (Sigma #P3803-100ML) extractions were performed and DNA was recovered by ethanol precipitation. Nested PCR was performed using the Herculase II Fusion DNA Polymerase (Agilent, #600679) and the DNA oligos indicated in **Table S1**. PCR1 products were purified using QIAquick PCR Purification kit (Qiagen #28104). PCR2 products were purified using Agencourt AMPure XP Beads (Beckman Coulter Life Sciences #A63880). DNA concentration was determined using Qubit dsDNA HS Assay Kit (Thermo Fisher #Q32854) and adjusted to 2 nM prior to sequencing. NGS was performed using the NextSeq 500/550 High Output Kit v2.5 75 cycles (Illumina #20024906).

### Screen analysis

Reads were demultiplexed using bcl2fastq Conversion Software v2.20 (Illumina) and fastx_toolkit v0.0.13. Sequencing adapters were removed using cutadapt v1.9.1 (66). The reference library was built using bowtie2 v2.2.9 (67). Read mapping was performed with bowtie2 allowing 1 seed mismatch in --local mode and samtools v1.9 (68). Mapping analysis and gene selection were performed using MAGeCK v0.5.6, normalizing the data with default parameters. sgRNA and gene enrichment analyses are available in **Table S5**-**S6**, respectively and full MAGeCK output at https://github.com/Simon-LoriereLab/crispr_isg_sarscov2.

### Generation of multi-guide gene knockout cells

3 sgRNAs per gene were designed (**Table S2**). 10 pmol of NLS-Sp.Cas9-NLS (SpCas9) nuclease (Aldevron #9212) was combined with 30 pmol total synthetic sgRNA (10 pmol for each sgRNA) (Synthego) to form RNPs in 20 μL total volume with SE Buffer (Lonza #V5SC-1002). The reaction was incubated at room temperature for 10 min. 2×10^5^ cells per condition were pelleted by centrifugation at 100*g* for 3 min, resuspended in SE buffer and diluted to 2×10^4^ cells/μL. 5 μL of cell solution was added to the pre-formed RNP solution and gently mixed. Nucleofections were performed on a Lonza HT 384-well nucleofector system (Lonza #AAU-1001) using program CM-120. Immediately following nucleofection, each reaction was transferred to a 96-well plate containing 200 μL of DMEM 10% FBS (5×10^4^ cells per well). Two days post-nucleofection, DNA was extracted using DNA QuickExtract (Lucigen #QE09050). Cells were lysed in 50 μL of QuickExtract solution and incubated at 68°C for 15 min followed by 95°C for 10 min. Amplicons were generated by PCR amplification using NEBNext polymerase (NEB #M0541) or AmpliTaq Gold 360 polymerase (ThermoFisher #4398881) and the primers indicated in **Table S3**. PCR products were cleaned-up and analyzed by Sanger sequencing. Sanger data files and sgRNA target sequences were input into Inference of CRISPR Edits (ICE) analysis https://ice.synthego.com/#/ to determine editing efficiency and to quantify generated indels (69). Percentage of alleles edited is shown in **Table 1**(n=3).

### SARS-CoV-2, SARS-CoV and MERS-CoV infection assays

A549-ACE2 cells were infected by incubating the virus for 1h with the cells maintained in DMEM supplemented with 1 μg/ml TPCK-trypsin (Sigma #4370285). The viral input was then removed and cells were kept in DMEM supplemented with 2% FBS. For 293T-ACE2 cells, infections were performed without TPCK-trypsin. MERS-CoV and SARS-CoV infections were performed in DMEM supplemented with 2% FBS and cells were incubated 1h at 37°C 5% CO2. Viral inoculum was then removed and replaced by fresh DMEM supplemented with 2% FBS. All experiments involving infectious material were performed in Biosafety Level 3 facilities in compliance with Institut Pasteur’s guidelines and procedures.

### Yellow Fever Virus and Measles Virus infection assays

Cells were infected with YFV (at an MOI of 0.3) or MeV-GFP (MOI of 0.2) in DMEM without FBS for 2h in small volume of medium to enhance contacts with the inoculum and the cells. After 2h, the viral inoculum was replaced with fresh DMEM 10% FBS 1% P/S. FACS analysis were performed at 24h p.i. Cells were fixed and permeabilized using BD Cytofix/Cytoperm (Fisher Scientific, # 15747847) for 30 min on ice (all the following steps were performed on ice and centrifuged at 4°C) and then washed tree times with wash buffer. Cells infected with YFV were incubated with the pan-flavivirus anti-Env 4G2 antibody for 1h at 4°C and then with Alexa 488 anti-mouse IgG secondary antibodies (Thermo Fisher, #A28175) for 45 min at 4°C in the dark. Non-infected, antibody-stained samples served as controls for signal background. The number of cells infected with MeV-GFP were assessed with the GFP signal, using non-infected cells as controls. Data were acquired with an Attune NxT Acoustic Focusing Cytometer (Life technologies) and analyzed using FlowJo software.

### Hit validation

2.5×10^4^ A549-ACE2 KO cells were seeded in 96-well plates 18h before the experiment. Cells were treated with IFNα and infected as described above. At 72h post-infection, supernatants and cellular monolayers were harvested in order to perform qRT-PCR and plaque assay titration. Infectious supernatants were heat-inactivated at 80°C for 10 min. For intracellular RNA, cells were lysed in a mixture of Trizol Reagent (Invitrogen #15596018) and PBS at a ratio of 3:1. Total RNA was extracted using the Direct-zol 96 RNA kit (Zymo Research #R2056) or the Direct-zol RNA Miniprep kit (Zymo Research #R2050). For SARS-CoV-2 detection, qRT-PCR was performed either directly on the inactivated supernatants or on extracted RNA using the Luna Universal One-Step RT-qPCR Kit (NEB #E3005E) in a QuantStudio 6 thermocycler (Applied Biosystems) or in a StepOne Plus thermocycler (Applied Biosystems). The primers used are described in **Table S4**. Cycling conditions were the following: 10 min at 55°C, 1 min at 95°C and 40 cycles of 95°C for 10s and 60°C for 1 min. Results are expressed as genome copies/mL as the standard curve was performed by diluting a commercially available synthetic RNA with a known concentration (EURM-019, JRC). For SARS-CoV and MERS-CoV, qRT-PCR were performed using FAM-labelled probes (Eurogentech) and the Superscript III Platinum One-Step qRT-PCR System (Thermo Fisher Scientific, #11732020). The cycling conditions were the following: 20 min at 55°C, 3 min at 95°C and 50 cycles of 95°C for 15 s and 58°C for 30 s. The primers used are described in **Table S4**. Standard curves were performed using serial dilutions of RNA extracted from and SARS-CoV and MERS-CoV viral culture supernatants of known infectious titer. For plaque assay titration, VeroE6 cells were seeded in 24-well plates (105 cells per well) and infected with serial dilutions of infectious supernatant diluted in DMEM during 1h at 37°C. After infection, 0.1% agarose semi-solid overlays were added. At 72h post-infection, cells were fixed with Formalin 4% (Sigma #HT501128-4L) and plaques were visualized using crystal violet coloration. Time-course experiments were performed the same way except that supernatants and cellular monolayers were harvested at 0h, 2h, 4h, 6h, 8h, 10h and 24h post-infection.

### Overexpression assay

2×10^5^ 293T-ACE2 cells were seeded in a 24-well plate 18h before experiment. Cells were transfected with 500 ng of plasmids expressing HA-DAXX WT, HA-DAXX 15KR and HA-DAXXΔSIM plasmids, using Fugene 6 (Promega # E2691), following the manufacturer’s instructions. HA-NBR1 was used as negative control. After 24h cells were infected at the indicated MOI in DMEM 2% FBS. When indicated, cells were treated with 10 mM of remdesivir (MedChemExpress #HY-104077) at the time of infection. For flow cytometry analysis, cells were fixed with 4% formaldehyde and permeabilized in a PBS 1% BSA 0.025% saponin solution for 30 min prior to staining with corresponding antibodies for 1h at 4°C diluted in the permeabilization solution. Samples were acquired on a BD LSR Fortessa and analyzed using FlowJo. Total RNA was extracted using a RNeasy Mini kit and submitted to DNase treatment (Qiagen). RNA concentration and purity were evaluated by spectrophotometry (NanoDrop 2000c, ThermoFisher). In addition, 500 ng of RNA were reverse transcribed with both oligo dT and random primers, using a PrimeScript RT Reagent Kit (Takara Bio) in a 10 mL reaction. Real-time PCR reactions were performed in duplicate using Takyon ROX SYBR MasterMix blue dTTP (Eurogentec) on an Applied Biosystems QuantStudio 5 (ThermoFisher). Transcripts were quantified using the following program: 3 min at 95°C followed by 35 cycles of 15s at 95°C, 20s at 60°C, and 20s at 72°C. Values for each transcript were normalized to expression levels of RPL13A. The primers used are indicated in **Table S4**.

### Microscopy Immunolabeling and Imaging

293T-ACE2 cells were cultured and infected with SARS-CoV-2 as described above. When indicated, cells were treated with 50 μM of GRL0617 (MedChemExpress #HY-117043) at the time of infection. Cultures were rinsed with PBS and fixed with 4% paraformaldehyde (electronic microscopy grade; Alfa Aesar) in PBS for 10 min at room temperature, treated with 50 mM NH4Cl for 10 min, permeabilized with 0.5% Triton X-100 for 15 min, and blocked with 0.3% BSA for 10 min. Cells were incubated with primary and secondary antibodies for 1h and 30 min, respectively, in a moist chamber. Nuclei were labeled with Hoechst dye (Molecular Probes). Images were acquired using a LSM700 (Zeiss) confocal microscope equipped with a 63X objective or by Airyscan LSM800 (Zeiss). Image analysis and quantification was performed using ImageJ.

### Western blot

293T-ACE2 cells were transfected with the indicated plasmids or treated with the indicated concentrations of GRL0617; with 10 μM of Masitinib (MedChemExpress #HY-10209); or with 10μM of MG132 (SIGMA #M7449), an inhibitor of the proteasome and infected with SARS-CoV-2. Cell lysates were prepared using RIPA lysis and extraction buffer (ThermoFisher #89901). Protein concentration was determined using Bradford quantification. Proteins were denaturated using 4X Bolt LDS Sample Buffer (Invitrogen) and 10X Bolt Sample Reducing Agent (Invitrogen). 40 μg of proteins were denatured and loaded on 12% ProSieve gel and then subjected to electrophoresis. Gels were then transferred (1h, 90V) to Western blotting membranes, nitrocellulose (GE Healthcare #GE10600002) using Mini Trans-Blot Electrophoretic Transfer Cell (Biorad #1703930EDU). Membranes were blocked with 5% BSA in PBS (blocking buffer) and incubated with primary antibodies diluted in blocking buffer. Membranes were washed and incubated with secondary antibodies diluted in blocking buffer. Chemiluminescent acquisitions were performed on a Chemidoc™ MP Imager and analysed using Image Lab™ desktop software (Bio-Rad Laboratories).

### Flow cytometry

For flow cytometry analysis, all cells were fixed with 4% formaldehyde. For intracellular staining, cells were permeabilized in a PBS/1% BSA/0.025% saponin solution for 30 min prior to staining with corresponding primary antibodies for 1h at 4°C and then secondary antibodies for 45min at 4°C, diluted in the permeabilization solution. Acquisition was done with Fortessa Cytometer and analyses with FlowJo software (Treestar Inc., Oregon, USA).

### Single-cell RNAseq analysis

Single cell RNAseq analysis were performed in the BioTuring Browser Software (v2.8.42) developed by BioTuring, using a dataset made available by Liao *et al.* (43) (ID: GSE145926). All processing steps were done by BioTuring Browser (71). Cells with less than 200 genes and mitochondrial genes higher than 10% were excluded from the analysis.

### Statistical analysis

GraphPad Prism was used for statistical analyses. Linear models were computed using Rstudio.

## Acknowledgements

We thank the Cytometry Platform, Center for Technological Resources and Research, Institut Pasteur, for cell sorting experiments. This work was funded by the Institut Pasteur Coronavirus Task Force, CNRS (UMR 3569), the Labex IBEID (ANR-10-LABX-62-IBEID) and by the ANR (ANR-20-COVI-000, projects IDISCOVR to M.V. and Alpha-COV to S.N.). A.M.K. is supported by a grant of the French Ministry of Higher Education, Research and Innovation. G.M. is supported by a grant from the Agence nationale de recherches sur le sida et les hépatites virales (ANRS). We thank Michael Emerman, Daniel Marc and Ignacio Caballero-Posadas for helpful comments on the manuscript. Illustrative figures in this manuscript were created with BioRender.com.

## Contributions

F.R. designed the research project. F.R. and M.V. secured the funding for the study. A.M.K., S.M.A., A.H., N.A., S.N., G.M., D.Q.T., M.C., T.V. and F.R. performed and analyzed the *in vitro* experiments. F.P. produced the stocks of lentiviruses. J.C.S., J.O. and K.H. generated and validated KO cell lines. T.B. performed the single-cell RNAseq data analysis. S.M. and F.D. performed the SARS-CoV and MERS-CoV experiments. A.B. and E.S.L. performed the bio-informatic analyses of the CRISPR/Cas9 screen. M.O., T.B., O.S., N.J., S.N., S.V.D.W. and M.V. analyzed the data and supervised the project. A.M.K., N.J. and F.R. wrote the manuscript. All authors edited the manuscript.

## Competing Interests

J.C.S., J.O. and K.H. are employees and shareholders from Synthego Corporation.

## Correspondence and requests for materials

should be addressed to either M.V., N.J., S.N. or F.R.

## Data availability

Raw NGS data was deposited to the NCBI GEO portal and is accessible with the number GSE173418.

## Notes

### Summary of Updates

We now show mechanistic evidence that DAXX inhibits an early step (likely viral transcription) of the SARS-CoV-2 life cycle, acting as early as 8-10h post-infection (Fig.3 revised). Using overexpression of viral proteins and different inhibitors, we show that the PLpro domain of SARS-CoV-2 Nsp3 is responsible for the proteasomal degradation of DAXX (novel Fig. 6). We also show that DAXX restriction exhibits some level of specificity: SARS-CoV, but not the related MERS-CoV or two other RNA viruses (Measles Virus and Yellow Fever Virus), was sensitive to DAXX restriction (Fig. 2 revised).

