## Supplemental Info for "Identification of DAXX As A Restriction Factor Of SARS-CoV-2 Through A CRISPR/Cas9 Screen"

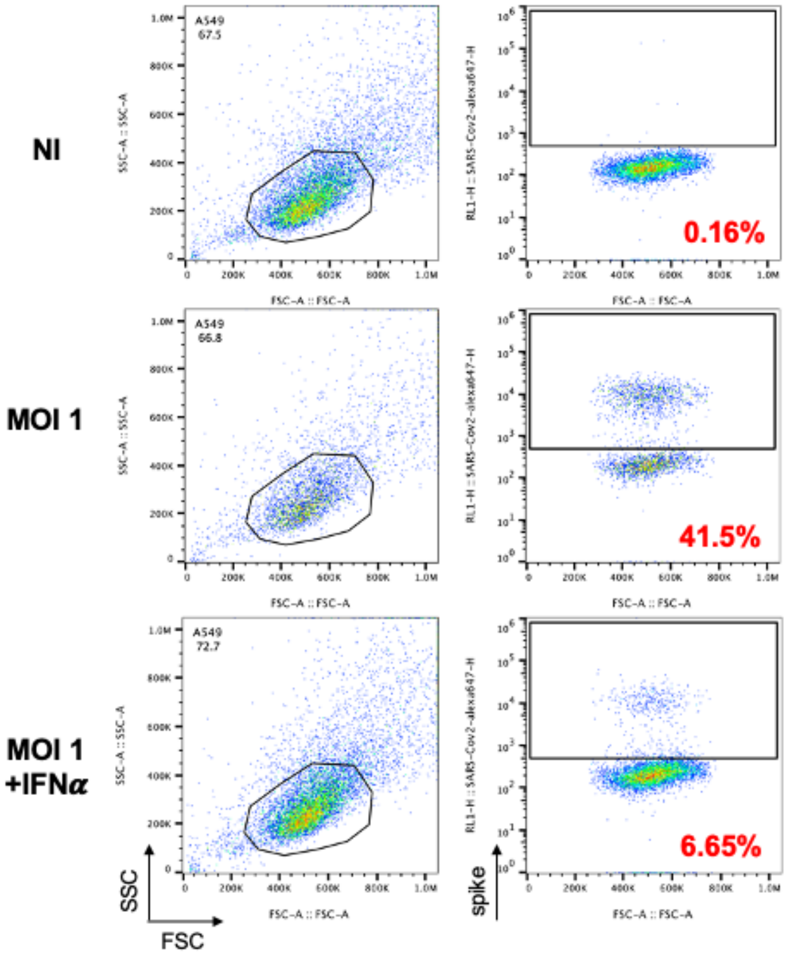


**Supplementary Figure 1: IFN-mediated restriction of SARS-CoV-2.** A549-ACE2 cells were pre-treated with 200 U/mL of IFNα and infected at MOI 1. Cells were labelled with anti-spike (S) antibody at 24h p.i. and the numbers of cells positive for S were analyzed by flow cytometry. Non-infected cells were used for gating controls. Percentage of infected cells are indicated in red. One representative experiment.

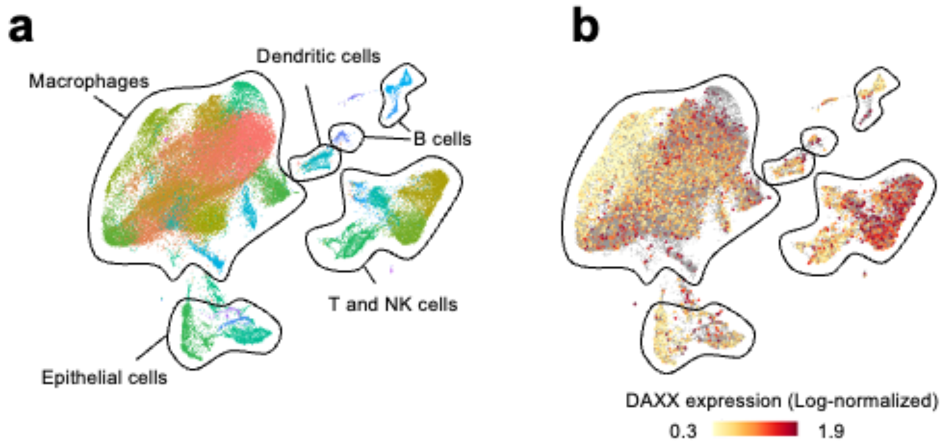


**Supplementary Figure 2: Expression of DAXX RNA in cells isolated from broncho-alveolar lavages. A:** Single cell RNAseq data of broncho-alveolar lavages from Liao *et al.* 2020 (43) (dataset ID: GSE145926) were analyzed using BBrowser Software. Colors indicate graph-based clusters. Cell types are indicated according to dataset metadata. Each dot represents an individual cell. **B:** Log-normalized expression of DAXX among single cells. Null values are excluded from the scale and indicated as grey dots.

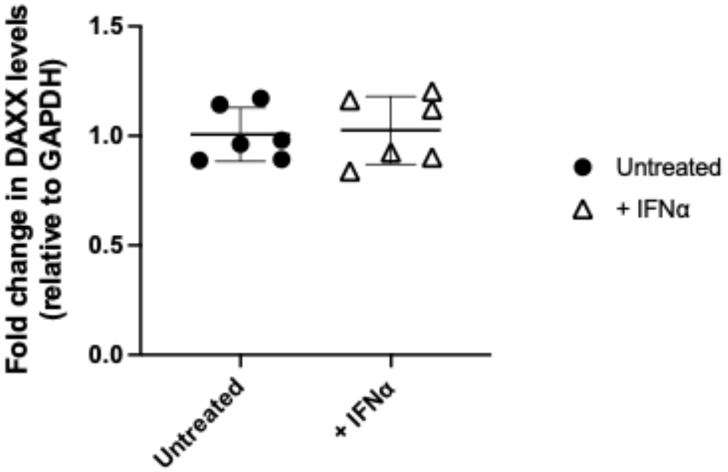


**Supplementary Figure 3: DAXX expression following IFNα treatment and SARS-CoV-2 infection.** A549-ACE2 WT cells were treated with 200 U/mL of IFNα for 24h in triplicates. Cell monolayers were harvested. Cellular RNAs were extracted and DAXX levels were quantified by qRT-PCR analysis. qRT-PCR against the housekeeping gene GAPDH was used as a control and to normalize DAXX levels. Fold changes in IFNα treated cells relative to control cells for 2 independent experiments are shown.

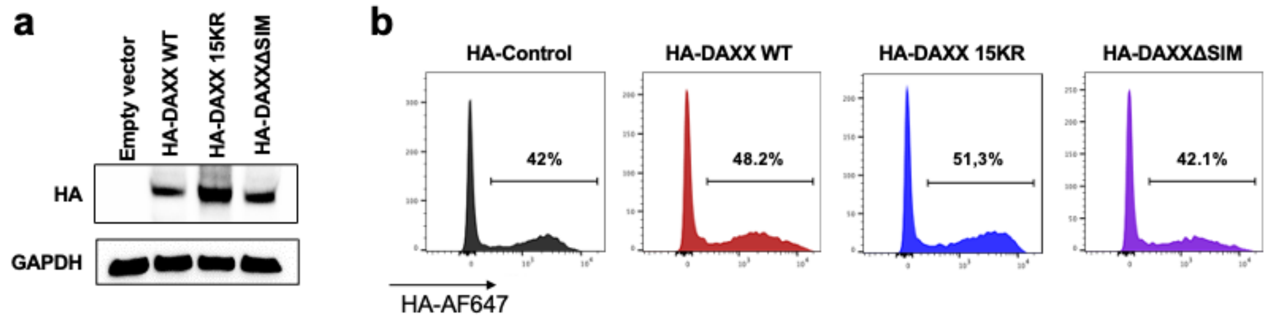


**Supplementary Figure 4: Expression of WT DAXX and mutants in transfected 293T-ACE2 cells.** 293T-ACE2 cells were transfected with the indicated HA-tagged DAXX constructs or with HA-NRB1 as a negative control. Levels of DAXX expression was measured by Western Blot (probing for HA ; GAPDH as a loading control) in (**a)** and by flow cytometry (intracellular HA staining) in (**b)**.

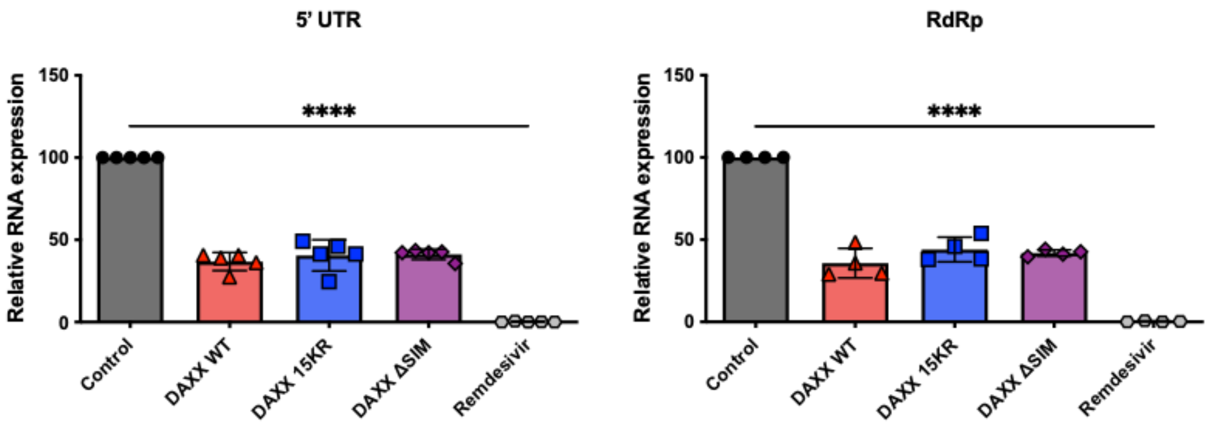


**Supplementary Figure 5: Effect of DAXX overexpression on SARS-CoV-2 transcription.** In parallel of the experiments shown in **Fig. 4d-e**, the intracellular levels of two viral transcripts (5' UTR ; RdRp) were quantified by qRT-PCR (normalized against RLP13a, ΔΔCt method). The mean of 4 independent experiments is shown. Statistics: one-way ANOVA: **** p-value < 0.0001.

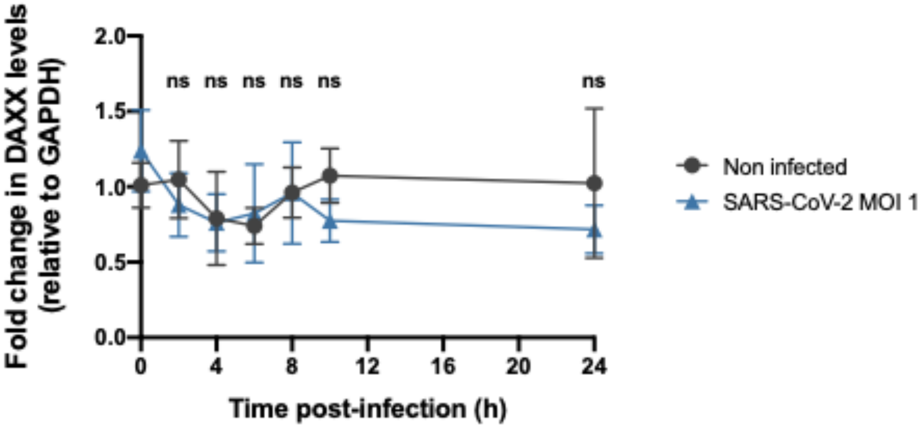


**Supplementary Figure 6: DAXX mRNA levels are not affected by SARS-CoV-2 replication.** A549-ACE2 WT cells were infected with SARS-CoV-2 at MOI 1. Cellular monolayers were harvested at the indicated time points and total RNA was extracted. The levels of DAXX RNA were determined by qRT-PCR analysis and normalized against GAPDH levels. The mean of 3 independent experiments performed in triplicates is shown. Statistics: 2-way ANOVA using Sidak's test. ns p-value > 0.05.

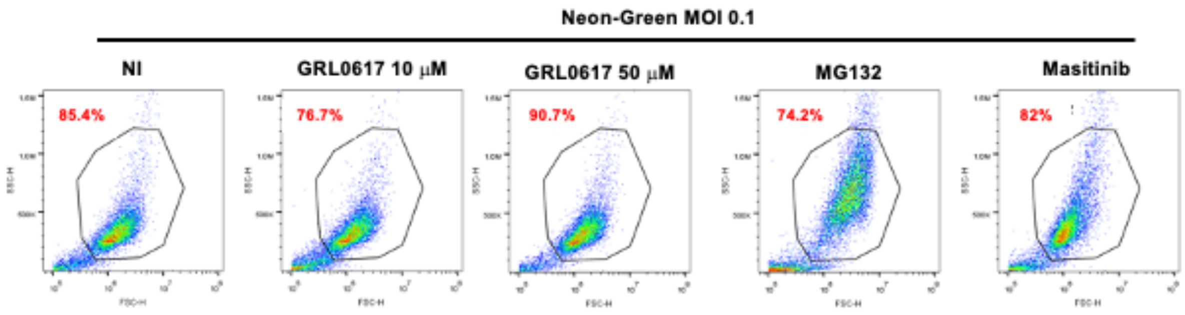


**Supplementary Figure 7: FACS analyses of 293T-ACE2 treated with inhibitors.**
The size (FSC) and granularity (SSC) of the cells used for the western-blots shown in **Fig. 6b** were evaluated by flow cytometry. The estimated percentage of live cells is indicated.

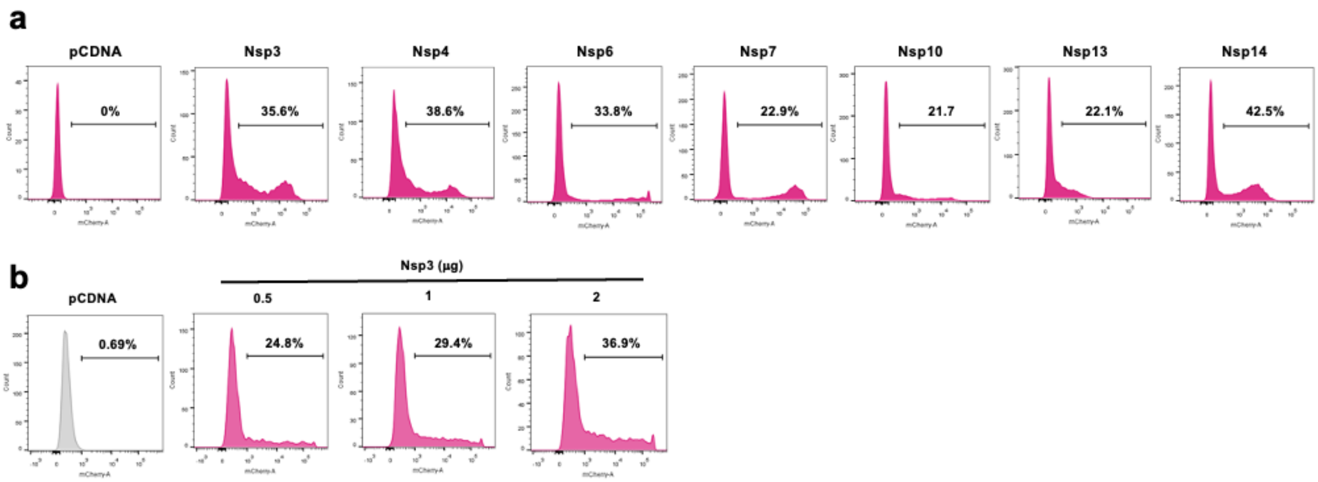


**Supplementary Figure 8: Expression levels of Nsp-mCherry fusion proteins in 293T-ACE2 cells.**

**(a)** The expression of SARS-CoV-2 Nsp proteins fused to mCherry (**Fig. 6d**) was evaluated by flow cytometry. **(b)** The expression of SARS-CoV-2 Nsp3-mCherry (**Fig. 6e**) was evaluated by flow cytometry.

**Table S1 : NGS oligos for the CRISPR/Cas9 screen.**

| Name | Sequence 5' ⇒ 3' |
|---|---|
| PCR1 Forward Primer | GAGGGCCTATTTCCCATGATTCCTTCA |
| PCR1 Reverse Primer | aacttctcggggactgtgg |
| PCR2 Forward Primer 1 (non infected cells) | AATGATACGGCGACCACCGAGATCTACACTCTTTCCCTACACGACGCTCTTCC GATCTAGTCAAtcttgtggaaaggacgaaacaccg |
| PCR2 Forward Primer 2 (infected cells) | AATGATACGGCGACCACCGAGATCTACACTCTTTCCCTACACGACGCTCTTCC GATCTATGTCAtcttgtggaaaggacgaaacaccg |
| PCR2 Reverse Primer | CAAGCAGAAGACGGCATACGAGATGTGACTGGAGTTCAGACGTGTGCTCTTCC GATCTtgccactttttcaagttgataacggact |

**Table S2: sgRNAs sequences for KO pool generation.**

| Gene | sgRNA1 | sgRNA2 | sgRNA3 |
|---|---|---|---|
| LY6E | GGCCUGGCACUCACCAAU GC | GCCGACCAUCUGCUCCGAC C | GGAGAAGCACAUCAGCGAG C |
| DAXX | CAGCACGAUGAUGCUGUU AG | CUCCCACCCACUCCCCAAU G | UCUGAGCCUCAUGGGGCCA G |
| APOL6 | UCCAGAGAUGACAGCAGU AG | CAAUAUCGUCUGCGAGGGC A | CUCAAAAAUAUUUUUUCUU C |
| HERC5 | CAACAACUGGGAGAGCCU UG | AGAAAAUUUCUAAAGCUUC U | CAGAUUAUCUUUGAGGCAG G |
| CTSL | UACUGUUGCCUCAUAUGG AU | AGGCUGCAAUGGUGGCCU AA | AGAUAAGCCUCCCAGUUUU C |
| IFI6 | GAAAAAGUGCUCGGAGAG CU | ACCUCCUCCGACGGCCAUG A | CCUCCAGGACUCGCAGUCG C |
| IFNAR1 | AAACACUUCUUCAUGGUA UG | GAGUGAAGAAAAGUUGCAU U | UUUACUUUAAAGAACUGGG A |

**Table S3: Primers used for sequencing of edited *loci.***

| Gene | Forward PCR primer | Reverse PCR primer | Sequencing primer |
|---|---|---|---|
| LY6E | CTGGCCACACTGTCTCAC TG | TGCATGGGAAATGAGG CTGT | CTGTCTCACTGTGTGTTTGAGT GTC |
| DAXX | GGAACTAGAAGGTTCAG GGGA | TGGAGGGGCTCATTCT GAGG | GAACTAGAAGGTTCAGGGGAA GAAGGAAG |
| APOL6 | TGTAGGGAGGTACAGGG AGG | TACCACTCACGATGCT GGTG | GATTCGAAGCTGAGAGTGGCA AGAATATC |
| HERC5 | GGAGGCTAGGTGAGAAG GGA | GTCTTTCCACTGAGAA GACAGGT | GAAGGGATGTAAACAGGGGTT TTAGAAAAC |

| | | | |
|---|---|---|---|
| CTSL | GGTAGACTTTAAAGTGATGTACAGTTCA | ACCCCACCCAGCCCTAATAT | CAGTTCACTTTTTAACAGTATTCAGATGTG |
| IFI6 | AGTAAAGAACGTCCCACCAGG | AAGTCCCTTCCCCTCTGTGA | AGTAAAGAACGTCCCACCAGG |
| IFNAR1 | AGAGTGGAAGGGTGTATGCT | CTTGGAAAGTGAACTGCTCTG | TGCTAAAATGTTAATAGGACATTAGCTCAA |

**Table S4: qRT-PCR primers and probes.**

| Targeted gene | Foward Primer | Reverse Primer | Probe |
|---|---|---|---|
| SARS-CoV-2 N | TAATCAGACAAGGAACTGATTA | CGAAGGTGTGACTTCCATG | |
| SARS-CoV-2 5' UTR | TGTCGTTGACAGGACACGAG | TTACCTTTCGGTCACACCCG | |
| SARS-CoV-2 RdRp | CATGTGTGGCGGTTCACTAT | TGCATTAACATTGGCCGTGA | |
| GAPDH | GAAGGTGAAGGTCGGAGTC | GAAGATGGTGATGGGATTTC | |
| DAXX | GGGCGACTATGTGAGCTGAA | GGCTTGTTGATGAGCCGCTC | |
| MERS-CoV E | GCA ACG CGC GAT TCA GTT | GCC TCT ACA CGG GAC CCA TA | CTC TTC ACA TAA TCG CCC CGA GCT CG |
| SARS-CoV N | TGG ACC CAC AGA TTC AAC TGA | GCT GTG AAC CAA GAC GCA GTA T | TAA CCA GAA TGG AGG ACG CAA TGG |
